# KAT: A K-mer Analysis Toolkit to quality control NGS datasets and genome assemblies

**DOI:** 10.1101/064733

**Authors:** Daniel Mapleson, Gonzalo Garcia Accinelli, George Kettleborough, Jonathan Wright, Bernardo J. Clavijo

**Affiliations:** Earlham Institute, Norwich Research Park, UK.

## Abstract

**Motivation:** *De novo* assembly of whole genome shotgun (WGS) next-generation sequencing (NGS) data benefits from high-quality input with high coverage. However, in practice, determining the quality and quantity of useful reads quickly and in a reference-free manner is not trivial. Gaining a better understanding of the WGS data, and how that data is utilised by assemblers, provides useful insights that can inform the assembly process and result in better assemblies.

**Results:** We present the K-mer Analysis Toolkit (KAT): a multi-purpose software toolkit for reference-free quality control (QC) of WGS reads and *de novo* genome assemblies, primarily via their k-mer frequencies and GC composition. KAT enables users to assess levels of errors, bias and contamination at various stages of the assembly process. In this paper we highlight KAT’s ability to provide valuable insights into assembly composition and quality of genome assemblies through pairwise comparison of k-mers present in both input reads and the assemblies.

**Availability:** KAT is available under the GPLv3 license at: https://github.com/TGAC/KAT.

**Supplementary Information:** Supplementary Information (SI) is available at Bioinformatics online. In addition, the software documentation is available online at: http://kat.readthedocs.io/en/latest/.

## 1 INTRODUCTION

Rapid analysis of high-throughput whole genome shotgun (WGS) datasets is challenging due to their large size (Metzker, 2010), with genome size and complexity creating additional challenges (Schatz *et al.*, 2012). Reference-free approaches for analysing WGS data typically involve examining base calling quality, read length, GC content (Yang *et al.*, 2013), and exploring k-mer (words of size *k*) spectra (Chor *et al.*, 2009; Lo and Chain, 2014). A frequently used reference-free quality control tool is FastQC (http://www.bioinformatics.babraham.ac.uk/projects/fastqc/)

K-mer spectra reveal information not only about the data quality (level of errors, sequencing biases, completeness of sequencing coverage and potential contamination) but also of genomic complexity (size, karyotype, levels of heterozygosity and repeat content) (Simpson, 2014). Additional information can be extracted through pairwise comparisons of WGS datasets (Anvar *et al.*, 2014), which can identify problematic samples by highlighting differences between spectra.

KAT, the K-mer Analysis Tookit, is a suite of tools for rapidly counting, comparing and analysing spectra for k-mers of arbitrary length directly from sequence data (see SI section 2 for a discussion on choice of *k* and SI section 3 for a comparison of k-mer tools).

## 2 THE K-MER ANALYSIS TOOLKIT

KAT is a C++11 application containing multiple tools, each of which exploits multi-core machines via multi-threading where possible. Core functionality is contained in a library designed to promote rapid development of new tools. Runtime and memory requirements depend on input data size, error and bias levels, and properties of the biological sample but as a rule of thumb, machines capable of *de novo* assembly of a dataset will be sufficient to run KAT on the dataset (see SI section 4 for details). K-mer counting in KAT is performed by an integrated and modified version of Jellyfish2 (Marçais and Kingsford, 2011), which supports large *k* values and is among the fastest k-mer counters available (Zhang *et al.*, 2014).

### 2.1 Assembly validation by comparison of read spectrum and assembly copy number

The KAT *comp* tool generates a matrix, with a sequence set’s k-mer frequency on one axis, and another’s set frequency on the other, with cells holding distinct k-mers counts at the given frequencies. When comparing reads against an assembly, KAT highlights properties of assembly composition and quality. If represented in a stacked histogram, read k-mer spectrum is split by copy number in the assembly (see SI section 5 for a primer on how to interpret KAT’s stacked histograms). In addition, KAT provides the *sect* tool necessary to study specific assembled sequences and track the k-mer coverage across both the read and the assembly spectra. This can help identify assembly artefacts such as collapsing or expanding events, or detect repeat regions. Figure 1 shows plots relating to two *Fraxinus excelsior* assemblies created from the same dataset using the *comp* and *sect* tools. The plots highlight different strategies taken by the assembler, in (a) and (c) we see some homozygous content being duplicated, and in (b) and (d) some heterozygous content eliminated.

**Fig. 1:**
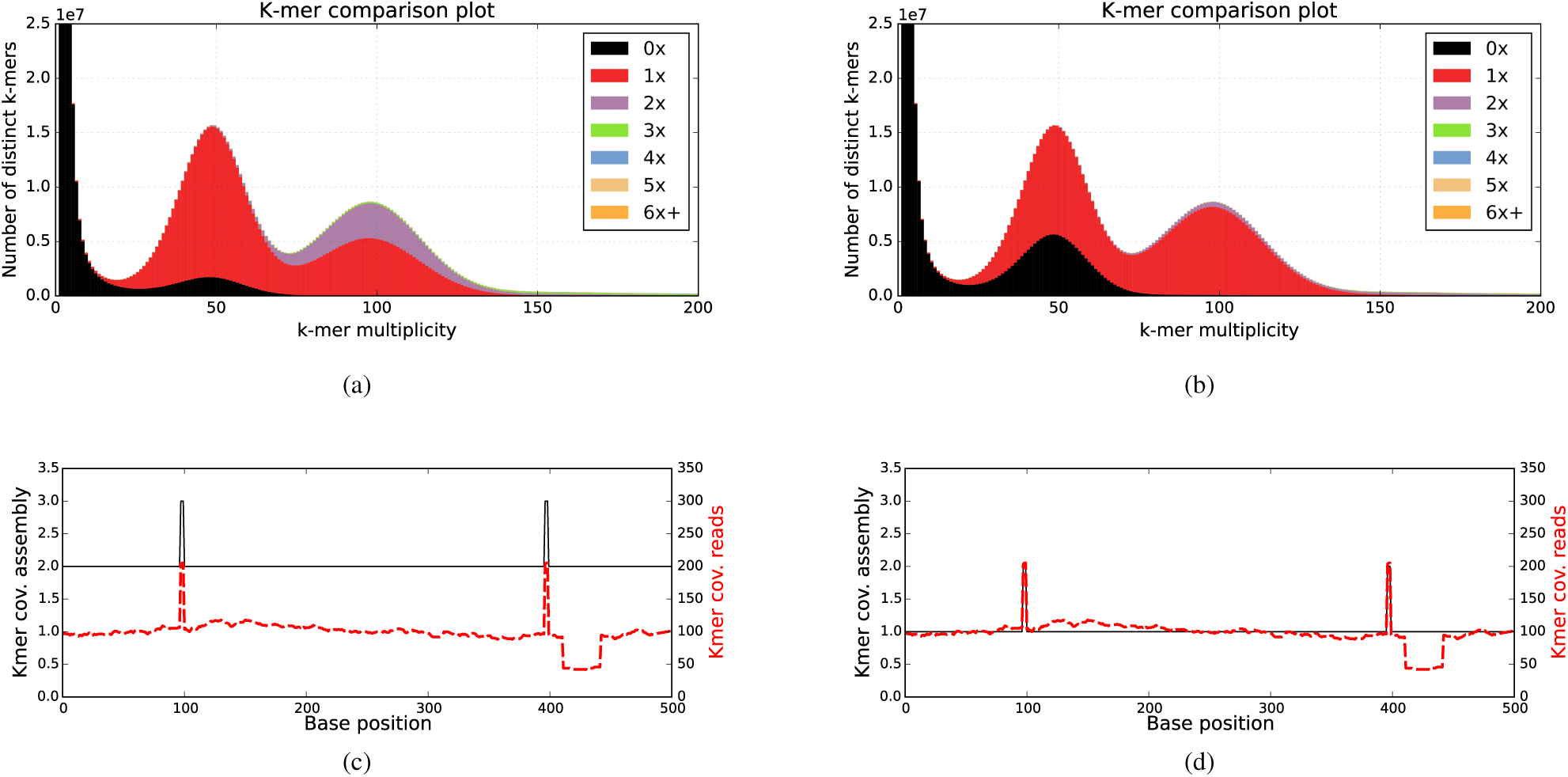
(a) and (b), generated using KAT *comp*, show read k-mer frequency vs assembly copy number stacked histograms for two different assemblies of a heterozygous *Fraxinus excelsior* genome http://ftp-oadb.tsl.ac.uk/fraxinus_excelsior. Read content in black is absent from the assembly, red occurs once, purple twice, etc. Both k-mer spectra show an error distribution under 25x, heterozygous content around 50X and homozygous content around 100X. (a) contains most (but not all) the heterozygous content, and introduces more duplications on homozygous content. (b) is more collapsed, including mostly a single copy of the homozygous content and less of the heterozygous content. (c) and (d), generated using KAT *sect*, show kmer coverage across example assembled loci. The assembly k-mer coverage (black line) of assembly (a) in plot (c) shows that the assembly has two copies of this locus, whereas the read k-mer coverage (red line) implies there should be only a single copy. This incorrect duplication has been corrected in assembly (b) with the read and assembly k-mer coverage agreeing in plot (d). The increased read and assembly k-mer coverage at positions 100 and 400 indicates small regions of repetitive sequence in the genome. The halved read k-mer coverage after position 400 indicates a heterozygous locus, which likely caused the duplication of this locus in the assembly (a). See SI Section 5 for a more extensive analysis of all sequences from this loci and their impact on (a) and (b).

### 2.2 Other KAT tools

KAT also includes the *hist* tool for computing spectrum from a single sequence set, the *gcp* tool to analyse gc content against k-mer frequency. The *filter* tool can be used to isolate sequences from a set according to their k-mer coverage or gc content from a given spectrum (see SI section 1 for details on all the tools). These tools can be used for various tasks including contaminant detection and extraction both in raw reads and assemblies, analysis of the GC bias and consistency between paired end reads and several libraries.

## 3 SUMMARY

KAT is a user-friendly, scalable toolkit for rapidly counting, comparing and analysing k-mers from various data sources. The tools in KAT assist the user with a wide range of tasks including error profiling, assessing sequencing bias and identifying contaminants and *de novo* genome assembly QC and validation.

## ACKNOWLEDGEMENTS

Thanks to David Swarbreck and Federica Di Palma for their support and all KAT users for their valuable feedback. This research was supported in part by the NBIP Computing infrastructure for Science (CiS) group.

